# Phenotypic plasticity, heritability, and genotype-by-environment interactions in an insect dispersal polymorphism

**DOI:** 10.1101/2025.03.18.643873

**Authors:** Lilian Cabon, Mahendra Varma, Gabe Winter, Anne Ebeling, Holger Schielzeth

**Author notes:** **Address for correspondence:** Lilian Cabon, Population Ecology Group, Institute of Ecology and Evolution, Friedrich Schiller University Jena, Dornburger Straße 159, 07743 Jena, Germany, Holger Schielzeth, Population Ecology Group, Institute of Ecology and Evolution, Friedrich Schiller University Jena, Dornburger Straße 159, 07743 Jena, Germany.

## Abstract

Evolutionary fitness is determined by the match between an organism’s phenotype and its local environment. When mismatched, individuals may disperse to more suitable habitats. For flightless insects, however, the range of dispersal is typically limited. Numerous flightless species have, therefore, evolved a dispersal dimorphism, that is, some individuals in otherwise short-winged populations develop long wings. Wing development may be genetically or environmentally determined, but these two drivers have rarely been analysed together. We studied the inheritance and density-dependent plasticity in the dispersal dimorphism of the meadow grasshopper *Pseudochorthippus parallelus*. Using a full-sib half-sib breeding design, we found that the development of long wings strongly depends on rearing density, with tactile stimulation being the most likely proximate cause. Additionally, we found heritable variation in the development of long wings, both in the propensity to produce long wings and in response to density (genotype-by-environment interactions). While at high and low densities, the environmental effect dominates, genetic variation takes effect mostly at intermediate densities. Our results have implications for the phenotype-environment match and ultimately the evolution of individualized niches. Induced dimorphisms represent a form of niche conformance (or adaptive phenotypic plasticity) and both genetic and induced dispersal dimorphisms facilitate niche choice in allowing individuals to sample a greater range of environments. Our study shows that niche-related polymorphisms can evolve via selection on the sensitivity threshold.

## Introduction

Evolutionary fitness is maximized when phenotypes match the local environment (Cody, 1974). Phenotypes are usually well adapted to their natural habitat (Ghalambor et al., 2007), but sometimes local conditions suddenly deteriorate, leading to poor phenotype-environment matches. Individuals may then be forced to disperse. Dispersal is a critical process in ecology that plays a key role in shaping range shifts, population dynamics, gene flow, and community structure (Mazzi and Dorn, 2012). Through dispersal, individuals have the opportunity to sample different environments and choose the environment that best matches their individual phenotype, a process known as niche choice (Trappes et al., 2022; Edelaar and Bolnick, 2019). Dispersal also has evolutionary consequences in maintaining genetic diversity within populations, promoting outbreeding, and facilitating adaptation to changing environmental conditions (Elbroch et al., 2009; Reuter et al., 2007; Pease et al., 1989). Despite the obvious beneficial consequences of dispersal, it poses a challenge for individuals with limited mobility, such as flightless insects.

Many predominantly flightless insects have evolved a dispersal dimorphism (Zera and Denno, 1997; Roff, 1986a; Harrison, 1980). This means that some individuals can become long-winged and dispersive in otherwise short-winged – and therefore flightless – populations (Zera and Denno, 1997). The switch between short- and long-winged phenotypes may be genetically determined (polymorphism in the strict sense) or be induced by the environment (often referred to as polyphenisms) (Zera and Denno, 1997). Most often the casual factors are unknown or mixed (as we show below), and we prefer to use the term polymorphism in the neutral sense to represent the phenotypic phenomenon of co-occurrence of discrete phenotypes within populations.

If developmental switches are induced by the environment, this represents a case of (adaptive) phenotypic plasticity. In the context of niche-related processes this can be viewed as a case of niche conformance, in which individuals adjust their phenotype to improve the phenotype-environment match (Trappes et al., 2022). At the same time, the development of long wings is also a facilitator of niche choice. While short-winged individuals have a limited range of environments to choose from, dispersing individuals have a substantially larger range of options (Takola and Schielzeth, 2022). The facultative development of long wings, thus, plays a dual role in affecting the phenotype-environment match and represents or facilitates the two niche-related mechanisms of conformance and choice.

While the ability for insects to fly seems advantageous, it often incurs physiological costs, such as reduced fecundity in females, reduced sperm production in males, shorter reproductive periods, and/or reduced longevity (Tigreros and Davidowitz, 2019; Vande Velde et al., 2012; Zhang et al., 2009; Roff, 1986a). This suggests that there might be a trade-off between flight capacity and reproductive outputs (although not supported for all species: (Jiang et al., 2010; Tigreros and Davidowitz, 2019). Additional costs can also arise from the increased risk of predation during dispersal (Zera and Denno, 1997), and uncertainty about finding suitable habitats. Accordingly, individuals should develop long wings only when the benefits of dispersion counterbalance these costs. This might be the case when environments change or fluctuate in such a way that individuals face poor local conditions, but there is a chance to reach more favourable patches within dispersal distance. The environmental factors affecting local conditions may be external (such as climatic factors) or be the results of the population dynamics of a species.

Indeed, wing dimorphism in insects is known to be influenced by population density (Lin et al., 2018; Poniatowski and Fartmann, 2009; Matsumura, 1996; Denno et al., 1991). An increase in population density – and, therefore, competition – results in a greater proportion of long-winged individuals in several species (Orthoptera, Tettigoniidae: Poniatowski and Fartmann, 2009; Hemiptera, Delphacidae: Denno et al., 1991, Lin et al., 2018, Matsumura, 1996; Hemiptera, Aphididae: Johnson, 1965, Sutherland, 1969). The presence of long-winged individuals in a population can mitigate the effect of elevated intra-specific competition, as long-winged individuals are generally considered the dispersive morph (Endo, 2006; Denno et al., 1991). Wing development is actually part of a larger dispersal syndrome that involves morphological, physiological and behavioural changes (Saastamoinen et al., 2018; Clobert et al., 2009). While long wings alone are not sufficient for dispersal to occur, they are an essential precondition and short-winged individuals certainly have low dispersal potential.

Wing dimorphism is not only influenced by environmental drivers, such as population density, but can also be genetically inherited. Wing dimorphisms appear to be controlled by polygenic inheritance in Orthoptera, Dermaptera and Hemiptera (Zera and Denno, 1997). Polygenic inheritance implies that many genetic loci (each typically of small effect) contribute to a phenotype. Although this usually produces continuous variation in phenotypes, it can also result in discrete phenotypes if there are thresholds for developmental switch points (Zhang et al., 2023; Saastamoinen et al., 2018; Roff and Fairbairn, 2007; Matsumura, 1996). Both genetic variation and environmental factors can influence whether an individual will take the long- or the short-winged developmental trajectory. Previous studies found the heritability of such threshold response in wing dimorphism to be between *h*^2^ = 0.52 and *h*^2^ = 0.72 in two species of crickets (Gryllidae) (Roff, 1990; Mousseau and Roff, 1989; Roff, 1986a). Since wing dimorphism is heritable and in face of the well-documented density dependency of wing dimorphisms, this raises interest in how much of the variance can be attributed to genetic versus environmental factors. Furthermore, there may be genetic variation in sensitivity to the environment (genotype-by-environment interactions, G x E). Studies that jointly estimated environmental and genetic drivers of wing dimorphisms are scarce in insects (Matsumura, 1996; Zhang et al., 2023), and, for Orthoptera, none have partitioned variance between those drivers nor accounted for genotype-by-environment interactions.

Here, we study the role of density-dependent phenotypic plasticity and inheritance in the wing dimorphism of the meadow grasshopper *Pseudochorthippus parallelus* (Orthoptera, Acrididae). This species inhabits a large range of grasslands across much of Europe (Chakrabarty and Schielzeth, 2020). Individuals are typically short-winged with both fore and hind wings being reduced (Figure 1). Fore wings in short-winged males are developed for stridulation (not flight), but are still shorter than fore wings of long-winged males (Köhler et al., 2017; Manzke, 1995). A small proportion of long-winged individuals (both females and males) can be found in natural populations (usually below 5%, personal observation). We used a full-sib half-sib breeding design to study the inheritance pattern of wing dimorphism, while raising the offspring in the lab at different densities to evaluate density-dependent patterns. We found that the proportion of long-winged individuals strongly increased with rearing density, but that families differed in the ratio of wing morphs in offspring, illustrating a genetic basis for the wing dimorphism. We also found G x E in response to density and total genetic variation had the largest phenotypic effect at intermediate densities. This illustrates that both environmental and genetic components are important, and that the dispersal dimorphism can evolve by natural selection.

**Figure 1:**
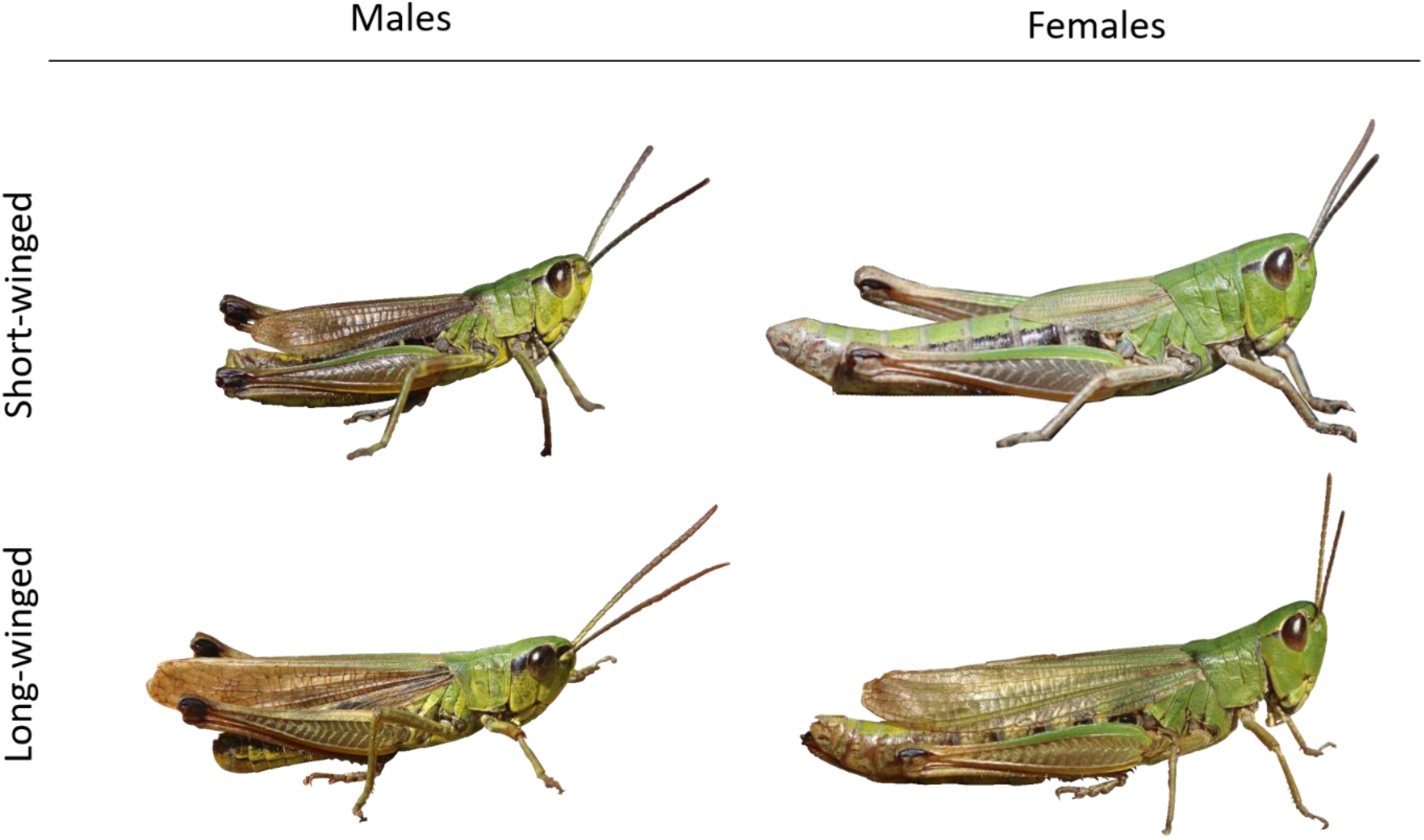
Wing dimorphism in the meadow grasshopper *Pseudochorthippus parallelus*. The dimorphism is defined by the length of the hindwings (which are rudimentary in short-winged individuals of both sexes), but it is visible also from the length of the forewings. Short-winged females have very short hind wings and fore wings, while fore wings of short-winged males can reach the tip of the abdomen. Long-winged individuals have fore wings exceeding the abdomen tip in males and reaching near or beyond the abdomen tip in females. Hind wings are of the same length as fore wings in long-winged individuals, but are clearly shorter in short-winged individuals.

## Materials and methods

### Parental generation

We set up a full-sib half-sib breeding design with meadow grasshoppers *Pseudochorthippus parallelus*. Parental individuals were caught from the field in the surroundings of Jena (50.94°N, 11.61°E) in June and July 2021. Only nymphal females were caught to ensure virginity. Males were mostly caught as nymphs, but a few adult males were also sampled. All parental individuals were short-winged, except for two long-winged males (0.8% of all parents). Parental individuals were housed in the laboratory in same-sex cages until maturity. A total of 172 mature females were then assigned to individual mating cages (22 x 16 x 16 cm), where they were maintained with freshly cut grass potted in small vials filled with water and a water tube for moisture. Grass pots were replaced three times per week. A small pot containing a 50:50 vermiculite-sand mixture was provided for egg deposition. A total of 68 males were mated to 1-4 females each by moving males every 2-3 days between mating cages, while ensuring females mated with only one male each. Males were kept in rotation and females were allowed to lay eggs until they died (or when breeding was terminated in early September 2021).

Sand pots were sieved once per week for the collection of egg pods. Egg pods are solid structures of 1-2 cm in length containing up to 12 eggs, and are typically buried in the sand. Egg pods were retrieved and placed in petri dishes lined with moist filter paper, while carefully documenting the cage of origin. All egg pods from the same cage and collection date were gathered on the same dish. A total of 1,560 egg pods were collected from 167 parental cages. In October, we replaced the filter paper with a 50:50 vermiculite-sand mixture and transferred petri dishes to refrigerators (approx. 5°C) for diapause. Egg pods were sprayed twice with a fungicide to prevent fungal growth. They were sprayed with water every week before diapause, for 6 weeks, while being kept at room temperature. They were then sprayed only biweekly during diapause, while being kept in the refrigerator.

### Offspring generation

A single offspring generation was raised to adulthood in five cohorts (cohorts represented the same generation, but had to be raised in separate time spans due to constraints in housing capacities). Individuals that hatched from the same petri dish were released together into a single offspring cage (22 x 16 x 16 cm). As different petri dishes produced different numbers of offspring, the densities per cage were the result of the hatching success in each petri dish. However, some densities were experimentally manipulated. After the large hatching peak had passed, cages hosting more than 8 offspring were evenly split into two cages (hereinafter referred as *receiving cages* as opposed to the *source cage* from which they originated). For logistic reasons, 32 cages (3% of the cages with more than 8 offspring) were not split and remained with densities ranging from 9 to 12 offspring per cage. Due to the large number of cages involved, it was operationally unfeasible to experimentally manipulate all offspring cage densities. Therefore, cages with up to 8 offspring were always left unsplit.

Some egg pods unexpectedly hatched before diapause on October 2021. Pre-diapause hatching came at a surprise and had not yet been documented for the meadow grasshopper. Offspring that hatched before diapause were raised as cohort 1. For this cohort, all egg pods collected from a parental cage on the same day were in a single petri dish. This happened because individual egg pods were isolated only after diapause to minimize the number of dishes to be tended during diapause. Therefore, for cohort 1, it is uncertain if hatchlings in an offspring cage came from the same or different egg pods of the same female. After diapause, each egg pod was placed in a Petri dish. For cohorts 2 to 5 (hatched in April, May, June, and July 2022, respectively), the offspring raised in one cage came from the same egg pod (with few exceptions). We emphasize that cohorts 1-5 represent a single offspring generation, and the separation into cohorts was done solely to produce a manageable number of active cages (at times, more than 400 offspring cages were maintained simultaneously).

Nymphs were maintained with *ad libitum* access to freshly cut grass potted in small vials filled with water and a water tube for moisture. Dead nymphs were recorded and removed. In total, 6,598 nymphs hatched, 1,683 (25%) nymphs died, 401 (6%) nymphs were lost for other reasons and 4,436 (67%) reached the imago stage about four weeks after hatching. Average full-sib family size of successfully developing offspring was 30.9 ± 20.9 individuals (mean ± SD, range 1-95) with 7.9 ± 4.2 (range 1-18) cages per family and 4.6 ± 1.9 (range 1-10) nymphal density classes represented per family (see below and Table 1 for offspring sample sizes).

**Table 1:** Sample sizes for different nymphal density classes.

|  | Nymphal density |  |  |  |  |  |  |  |  |  |  |  |
| --- | --- | --- | --- | --- | --- | --- | --- | --- | --- | --- | --- | --- |
|  | 1 | 2 | 3 | 4 | 5 | 6 | 7 | 8 | 9 | 10 | 11 | 12 |
| Female offspring | 82 | 207 | 250 | 308 | 345 | 356 | 235 | 225 | 88 | 40 | 16 | 8 |
| Male offspring | 87 | 171 | 239 | 316 | 357 | 288 | 244 | 215 | 101 | 40 | 6 | 4 |
| Total offspring | 169 | 378 | 489 | 624 | 702 | 644 | 479 | 440 | 189 | 80 | 22 | 12 |
| Families | 102 | 99 | 92 | 89 | 81 | 62 | 43 | 37 | 18 | 8 | 2 | 1 |
| Cages | 169 | 191 | 164 | 158 | 142 | 108 | 69 | 55 | 21 | 8 | 2 | 1 |

Offspring densities were quantified as the number of offspring per cage. We refer to the number of hatchlings that were released into a cage as *hatchling density* and to the number of adults that developed in a cage (after early nymphal deaths and splitting of cages with more than 8 individuals) as *nymphal density*. Most nymphal mortality occurred within the first few days after hatching, such that hatchling density roughly represents density within the first week of nymphal development, while nymphal density mainly represents the density during weeks 2-4 of nymphal development (in which little mortality occurred). Nymphal density was analysed by including both source (unsplit) cages and receiving cages (with densities after splitting). Hatchling density did not include receiving cages (9% of all cages) and included source cages with their densities before splitting. Hatchling density, in principle, also represents the number of embryos that successfully emerged from a single egg pod. However, offspring from cohort 1 and 4.5% of the cages from cohorts 2 to 5 came from more than one egg pod and for those cases the number of developing embryos per egg pod was unknown. Since embryos may influence each other during development, we analysed *embryo density* as the number of hatchlings among the subset of cages that derived from a single egg pod.

These densities could be confounded with maternal fecundity. However, most of the fecundity of female grasshoppers is determined by the total number of egg pods produced rather than the number of fertilized eggs per egg pod (which is also, perhaps mostly, affected by male fertility (Butlin et al., 1987). The total number of offspring per female was correlated with offspring density per cage in our experiment, though the correlation was not strong (r = 0.33). While we cannot rule out an effect of maternal fecundity on density, we consider it more likely that any effects operated via embryo number (within egg pods) or nymphal density.

### Offspring phenotyping

After the final moult, mature offspring were collected from their offspring cages. We recorded offspring cage, sex and wing morph. Short-winged meadow grasshoppers have their hind wings rudimentarily developed. Fore wings of short-winged females reach the middle of the abdomen (Figure 1). Short-winged males have better developed forewings that can reach the tip of the abdomen. Fore and hind wings exceed the abdomen tip in long-winged males while almost reaching the abdomen tip in long-winged females. In some intermediate cases (2.5%), we checked the length of the hind wings and assigned to long-winged individuals the ones with well-developed hind wings.

### General statistical analyses

Before delving into the animal model analyses (see below), we estimated sex-biases in the probability of developing short or long wings. This was done by χ^2^ tests on pooled data. To further evaluate sex-differences in the wing length development across the environmental gradient, we used correlation analysis for the proportion of short-winged individuals for each density class on pooled data (females vs. males). Since there was no evidence of any sex-specific effect even in these analyses on pooled data (see results), we did not include offspring sex in the animal model analysis.

Furthermore, we evaluated the different measures of density as described above (nymphal density, hatchling density, embryo density) and also proportional survival per cage before fitting the animal model. This was done on pooled data in a generalized linear model with binomial error structure, logit link and density as the only predictor.

### Animal model analysis

We fitted animal models (Kruuk, 2004) to decompose the variance into environmental and genetic sources of variation using the R package brms version 2.20.4 (Bürkner, 2017). Our basic animal model is represented by the following phenotypic equation and population-level variances:

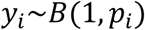

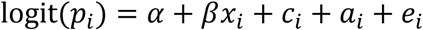

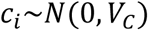

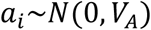

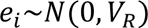

where y*i* represents an indicator of wing length of offspring *i* (y*i* = 1 for long-winged and y*i* = 0 for short-winged individuals) that is modelled by a Binomial distribution with probability *p*_i_ for offspring *i* (note that parameter n = 1 for binary data). Probability *p*_i_ is modelled on the logit scale where α represents the intercept, x*i* represents the density experienced by individual ***i*** (centred to zero at a density of 6.5, which represents the midpoint between densities 1 and 12), β represents the population-level slope of the density effect, c*i* represents the cage identity (permanent environment) random effect of individual ***i*** with a population-level variance *V*_*c*_, a*i* represents the additive genetic effect (individual identity linked to the two-generation pedigree) with a population-level additive-genetic variance *V*_*A*_ and e*i* represents the residual effect for individual ***i*** with a population-level residual variance *V*_*R*_. Assuming that all wild-caught individuals were unrelated, the additive genetic relatedness matrix used to estimate *V*_*A*_ consisted of values 0 for unrelated individuals, 0.5 for full-siblings and 0.25 for half-siblings. We refer to the variance *V*_*A*_ as the pedigree effect here (sometimes also referred to as the animal effect).

Furthermore, we fitted a random-slope animal model that estimates the additive-genetic variance of response to nymphal density (as well as the random intercept-slope correlation). The random-slope animal model is represented by the following phenotypic equation and population-level variances:

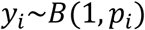

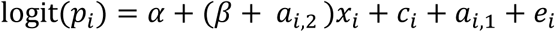

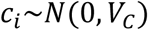

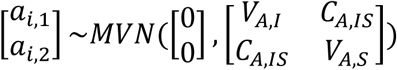

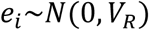

where *a*_*i*,1_ now represents the additive genetic effect with the population-level additive-genetic random intercept variance *V*_*A*,*I*_, *a*_*i*,2_ represents the additive genetic effect for the response to density with the population-level additive-genetic random slope variance *V*_*A*,*S*_, and *c*_*A*,*IS*_ represents the population-level additive genetic intercept-slope covariance. Other terms are the same as above.

We used default minimally informative priors for all parameters estimated by the model. The (non-)informativeness of the priors was checked with posterior predictive checks. For the fixed effect slope β the default prior is an improper flat prior. For random effects variances *V*_*c*_, *V*_*A*_, *V*_*A*,*I*_ and *V*_*A*,*S*_ default priors are half student-t priors on the respective standard deviation with 3 degrees of freedom and a scale parameter adjusted to the standard deviation of the response on the link scale (http://paul-buerkner.github.io/brms/reference/set_prior.html). Finally, for the additive genetic variance-covariance matrix there is a prior for the Cholesky factor of the correlation matrix with η = 1. We ran two chains of the Bayesian model with a warmup of 50,000 iterations followed by 120,000 iterations for sampling from the posterior distributions keeping every 70^th^ sample. Convergence was checked from trace plots, by *R*^ criterion (all *R*^ ≤ 1.008) and posterior effective sample size (all ESS ≥ 1,774, out of 2,000 posterior samples across the two chains). All analyses were done in R 4.1.3 (R Core Team, 2022).

Evaluation of the statistical significance of variance components can be difficult in Bayesian models, since variances are bound at zero and posterior medians and credibility intervals are often larger than zero even in the absence of a statistically significant effect (Pick et al., 2023). Since the statistical significance of the G x E effect (estimated in the form of random-slope variation) is of particular interest in our analysis, we used simulations to evaluate the posterior distribution in comparison to a null-model reference scenario. This was done by randomizing the vector of densities and fitting this randomized vector as an additional fixed effect and random-slope term – instead of the random slope term for the actual density data. All other terms in the model remained the same, that is, we fitted a population-level slope for actual density data and individual identity linked to the pedigree as a random (intercept) effect. We compared the posterior distribution for the random slope variance for the real-data model to the posterior distribution for the randomized random-slope term for 19 iterations of randomization. We chose 19 iterations of randomization because our observed data represents one particular randomization for which we aimed to determine whether its distribution could have risen by chance or represents a deviation from the null hypothesis represented by the 19 randomizations. Therefore, if the observed distribution represents an actual deviation from the null hypothesis, we can conclude to a deviation with a confidence level of 95% (19/20).

Variance components were calculated following (Nakagawa and Schielzeth, 2010) with the distribution-specific variance of the binomial distribution approximated by π*2/3 and the residual variance fixed to 1. These variances give the link-scale variances. We can also calculate the data-level variances (following Nakagawa and Schielzeth, 2010), Table 1, using the average proportion of long-winged individuals for *P*), which, reassuringly, gave highly similar variance ratios. For random-slope models, however, variance components depend on the value of the covariate and we used a recently proposed method for conditional variance decomposition (Schielzeth and Nakagawa, 2022). The total (marginalized) additive genetic variance (arising from random intercept and random-slope additive genetic variation) is given by *V*_A,M_ = *V*_A,I_ + *V*_A,S_*V*_x_ + µ²V_A,S_ + 2 µ C_A,IS_, where *V*_A,I_, *V*_A,S_ and C_A,IS_ are as above (for the random-slope model), and µ and *V*_x_ the mean and variance of nymphal densities, respectively. The variance exclusively explained by random-slopes is *V*_s_ = *V*_A,S_*V*_x_ + µ²*V*_A,S_. Both, *V*_A,M_ and *V*_S_ can be set in relation to the total phenotypic variance.

## Results

### General patterns

We assessed wing morph phenotypes of 4,228 offspring (51.1% females, 48.9% males). Of these offspring, 45.6% were scored as short-winged and 54.4% as long-winged. The proportion of long-winged individuals was similar in both sexes (53.4% in females, 55.6% in males, χ^2^_1_ = 2.02, p = 0.16).

Wing morph strongly depended on the nymphal density, with 99.4% of offspring raised in cages with a single nymph being short-winged (only 1 individual out of 168 was long-winged) and 83.8% of offspring raised in cages with 10 nymphs being long-winged (67 out of 80 individuals). The increase in the proportion of long-winged individuals appeared rather linear with no clear density-dependent switch point (Figure 2). There might seem to be a slight reduction in the proportion of long-winged individuals at nymphal densities greater than 10 individuals, though with just two cages of density 11 and one cage of density 12, neither of these proportions was significantly different from the nymphal density of 10 individuals (respectively χ^2^_2_ = 0.15, p = 0.70; χ^2^_2_ = 0.11, p = 0.74). Wing morph across the range of nymphal density (12 density classes) was highly correlated between the sexes (r = 0.99, t_10_ = 18.7, p < 0.001).

**Figure 2:**
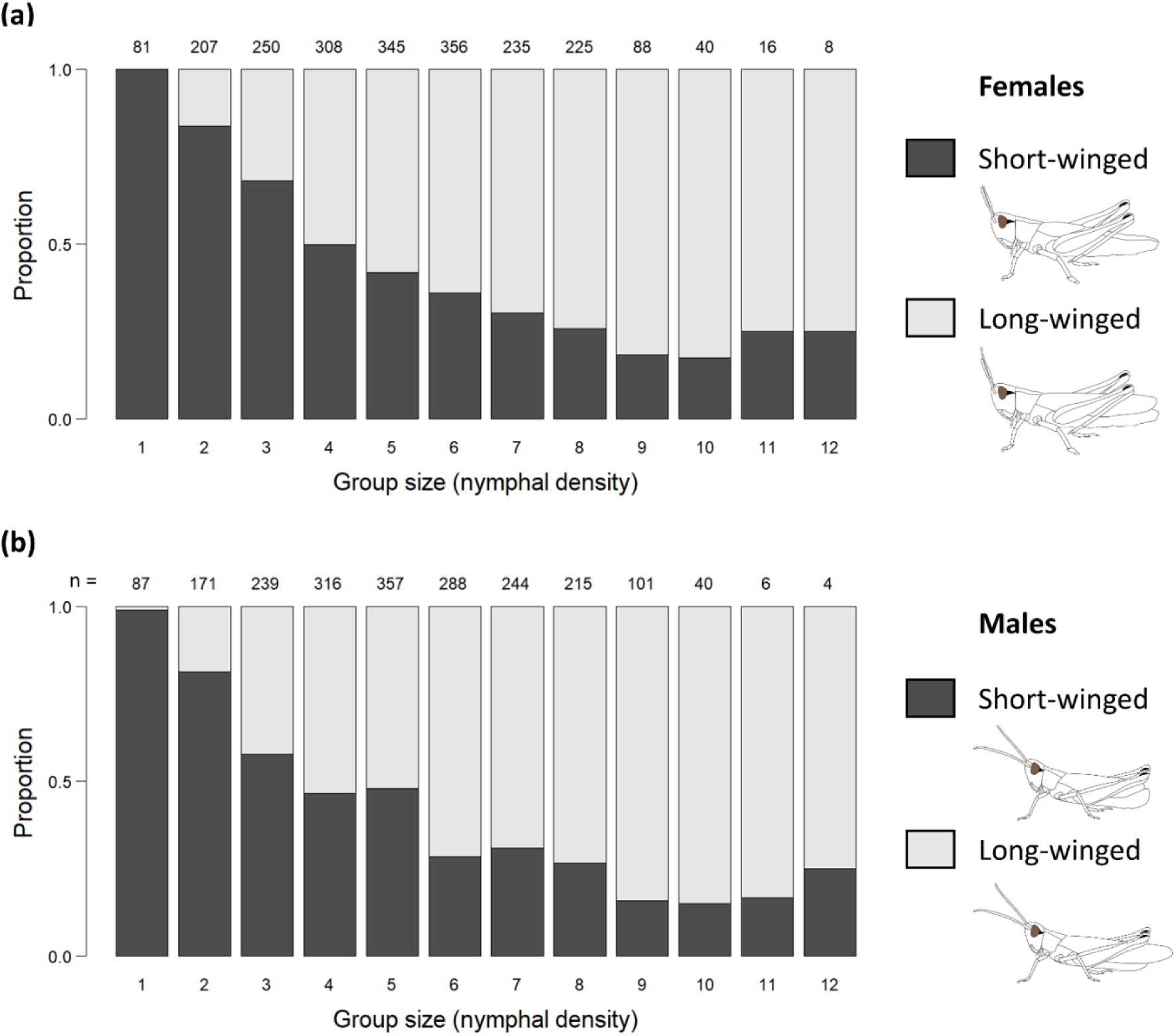
Density-dependence in wing development in (a) females and (b) males of the meadow grasshopper. Numbers above bars show the number of phenotyped individuals per density class.

Nymphal density was highly correlated to hatchling density (r = 0.67, t_993_ = 28.5, p <0.001) and to embryo density (r = 0.70, t_887_ = 29.1, p <0.001). It is therefore difficult to disentangle which aspect of density during development most strongly influence wing morph. Since datasets slightly differ between different measures, formal model comparisons are not possible. We, therefore, (informally) compared patterns using all cages where densities were not artificially altered (not moving in or out). A true causal predictor would show similar slopes in those three datasets. Nymphal density appeared to be better and more consistent predictor than hatchling density (Figure 3a,b). However, embryo densities appears to be equally good (Figure 3a,c). We also investigated whether the proportion of hatchlings surviving to adulthood – a potential proxy for the goodness of conditions during development – would predict offspring wing morph. However, cage-wise survival was a poor predictor of offspring wing morph (Figure 3d).

**Figure 3:**
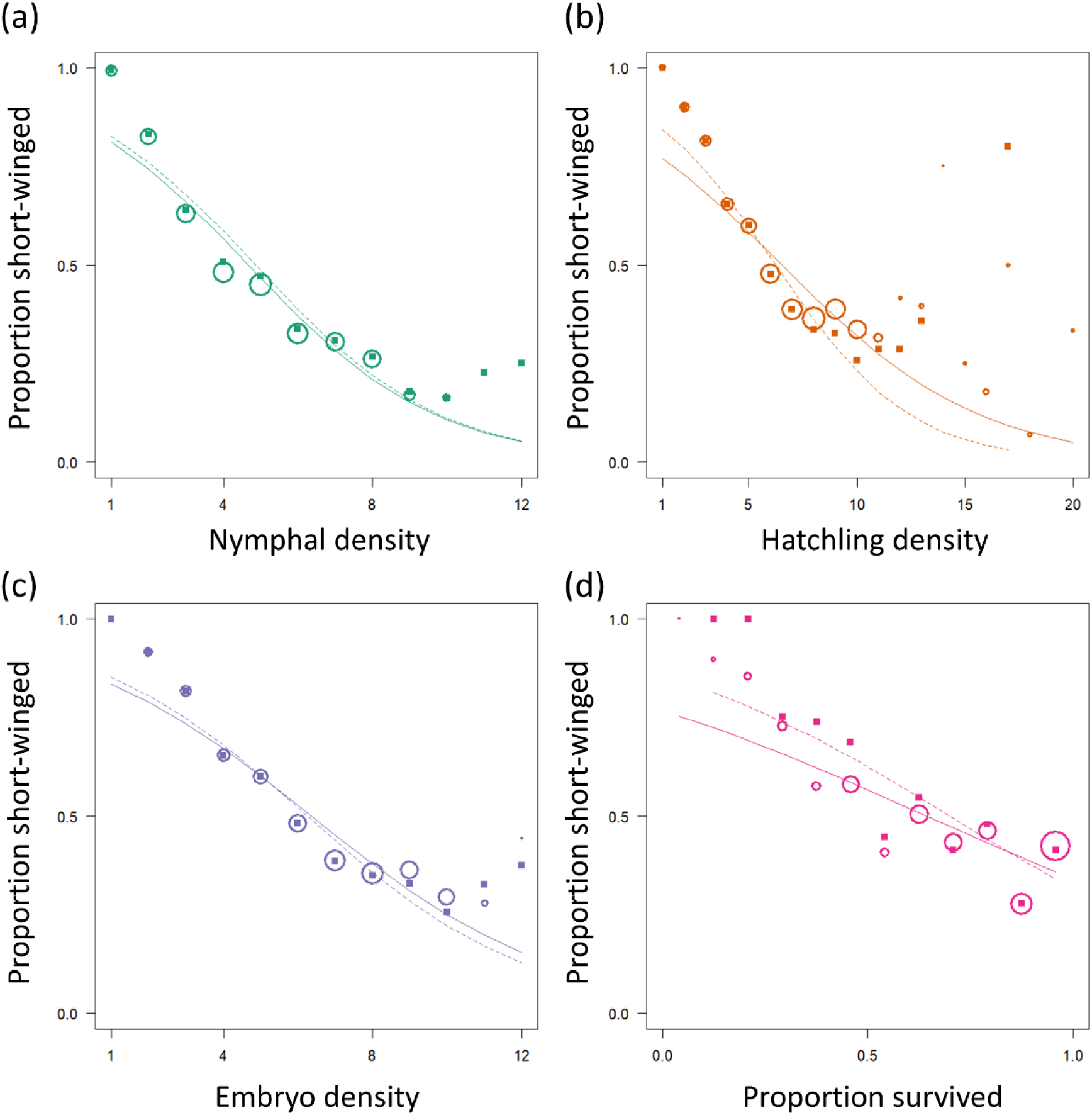
The effect of (a) nymphal density, (b) hatchling density, (c) embryo density, and (d) cage-wise survival on wing morph frequency in the meadow grasshopper *Pseudochorthippus parallelus*. Circles show the proportion of short-winged individuals per density class (with circle sizes proportional to the square-root sample size) and solid lines show binomial GLM fits for the pooled data with density as the only predictor. Solid squares show data based on only those cages that were never split or moved (excluding source and receiving cages) with dashed line showing the respective GLM fits.

### Animal model analysis

The animal model analysis demonstrated heritable genetic variation in offspring wing morph (Table 2). Density explained about 22.4% of the variance, while genetic relatedness (additive genetic variation) explained 18.9% and cage identity (shared early rearing effects, egg pod effects and potentially maternal effects) explained 7.2% of the variance. We then fitted a random-slope animal model that allowed the nymphal density slope to vary from additive genetic variation. The posterior distribution of the random-slope genetic variance was small, but shifted away from zero as compared to a randomized random-slope term (Figure 4c). The marginalized additive genetic variance increased to 19.1%, while additive genetic variation for the nymphal density slopes (that is, gene-by-environment interactions) explained 2.4% (Table 2, Figure 4a). The conditional genetic variance along the nymphal density gradient was small at very low densities (5.4 to 8.4% at nymphal density of 1 and 2, respectively), maximized at moderate densities (13.9% at a density of 4) and extremely low at high densities (less than 2% at densities of 9 or higher) (Figure 4b).

**Figure 4:**
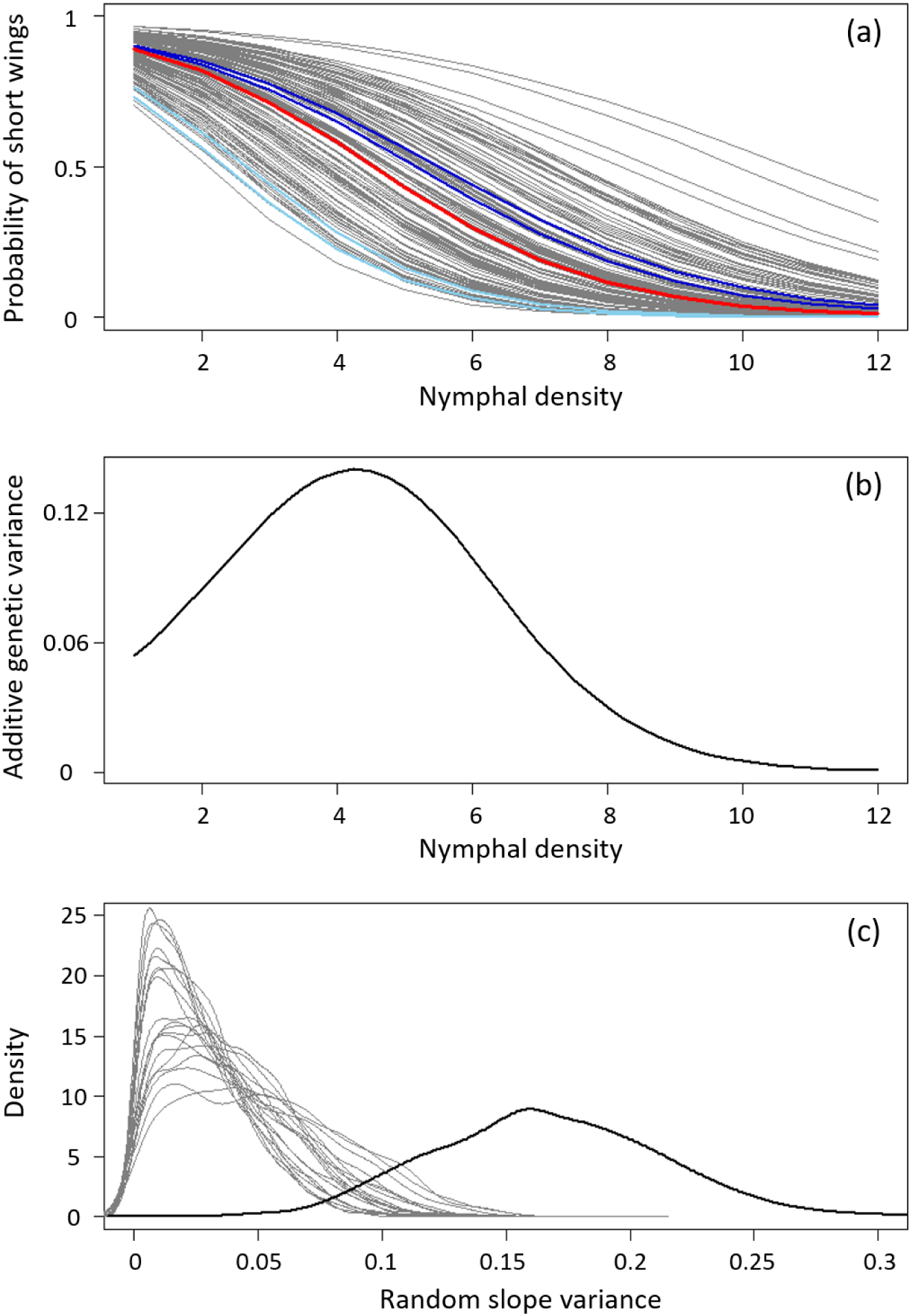
Effects of the family of origin on the propensity of offspring to develop short wings depending on the nymphal density. (a) Family-wise patterns in the probability of producing short-winged offspring depending on the nymphal density. Grey curves are family-wise patterns with both short-winged parents. Blue curves are family-wise patterns with a short-winged mother and a long-winged father (different shades of blue representing different fathers). The red curve is the population-wide pattern. (b) Variance explained by additive genetic effects in the development of short wings depending on the rearing density. (c) Comparison of the posterior distribution for the random-slope variance (bold) to 19 replicate null scenarios (posterior distribution for the random-slope variance based on randomized data, thin lines).

**Table 2:** Summary of the random-slope animal model analysis. Wing morph was modelled as a binary response (0 for short-winged, 1 for long-winged), nymphal density as a continuous fixed effect, cage identity as a blocked random effect and the pedigree effect was estimated by linking individual identity to the pedigree as a continuous random effect. Errors were modelled as binomial with the logit link function. Variance components are shown as proportions of the total phenotypic variance. HPD = Highest probability density.

| Fixed effects | Posterior median | Posterior SD | 95% HPD interval |
| --- | --- | --- | --- |
| Intercept | 1.146 | 0.168 | [0.835; 1.470] |
| Nymphal density slope | 0.589 | 0.038 | [0.517; 0.665] |
| Random effects | Posterior median | Posterior SD | 95% HPD interval |
| Pedigree | 1.485 | 0.314 | [0.982; 2.202] |
| Cage identity | 0.572 | 0.126 | [0.334; 0.807] |
| Pedigree x Density | 0.191 | 0.108 | [0.037; 0.429] |
| Cov(Cage identity, Pedigree x Density) | 0.183 | 0.077 | [0.056; 0.343] |
| Variance component | Posterior median | Posterior SD | 95% HPD interval |
| Nymphal density | 0.224 | 0.021 | [0.185; 0.266] |
| Pedigree | 0.189 | 0.031 | [0.139; 0.261] |
| Pedigree x Density | 0.024 | 0.013 | [0.005; 0.052] |
| Cage identity | 0.072 | 0.015 | [0.045; 0.101] |
| Residual | 0.513 | 0.030 | [0.450; 0.568] |
| Marginalized pedigree effect | 0.191 | 0.108 | [0.037; 0.429] |

The two long-winged founder males produced a total of 154 offspring with 5 different females. One full-sib family was too small (4 offspring) for family-wise analyses. One of the long-winged males produced clearly more long-winged offspring than the population average with two different females, while the other male produced rather average ratios with two different females (Figure 4a).

## Discussion

We analysed phenotypic plasticity and heritability of a dispersal dimorphism in the meadow grasshopper *Pseudochorthippus parallelus*. All but two founders of our breeding population were short-winged individuals, so that the predisposition to develop long wings might have appeared to be low. However, more than half of the 4,227 lab-reared offspring were long-winged. Nymphal density had the most pronounced influence on the development of long wings. Nearly all individuals in single-housed offspring were short-winged (>99%) and more than 80% were long-winged when housed in groups of 8 or more individuals. Besides the strong effect of nymphal density, we found significant heritable variation in the development of long wings. While about 22% of the variation was explained by offspring density another 19% were explained by additive genetic effects, and 2-3% by genotype-by-environment interactions.

Wing reduction is associated with habitat stability in several insect species (Zera and Denno, 1997; Denno et al., 1996; Roff, 1986b). The meadow grasshopper inhabits a broad range of grasslands that seem to be stable in time (Bridle et al., 2001) and, thus, fits this general pattern. Field populations consist almost entirely of short-winged individuals (rough estimation of 1 to 5% long-winged, personal observation). Densities were much higher in the laboratory (with 5.2-62.5 ind/m^2^, considering the surface of the six sides of the cage, as grasshoppers like to perch on all six surfaces when maintained in the lab) compared to natural populations (about 0.1 ind/m^2^, and exceptionally up to 3.16 ind/m^2^; Gardiner et al., 2002, Gardiner and Hill, 2006), implying that *V_A_* in the field is likely to be very low. This difference in densities is the most likely explanation for the large difference in the frequencies of long-winged individuals in the laboratory as opposed to natural populations. Such extreme cases in lab densities allowed us to explore patterns over a wide range of densities and depict that the environment is the main driver of long wing development at extreme densities. The observed density-dependence of the dispersal dimorphism is in line with the well-known density-dependent phase polyphenism in locusts (Song, 2011), and also with studies on various other Orthoptera (An et al., 2012; Poniatowski and Fartmann, 2009; Matsumura, 1996). Studies focusing on heritable components in crickets did not manipulate and assess rearing densities (Roff, 1990; Mousseau and Roff, 1989; Roff, 1986a), illustrating that studies usually focus on either the environmental or the genetic component.

The proximate cause of density-dependent dispersal dimorphism in Orthoptera may be, more generally, the quality of the local environment. Competition at high densities can affect food availability (Liu et al., 2007; Belovsky and Slade, 1995), mating opportunities (Lehmann and Lehmann, 2007), or disease transmission (Bailes et al., 2020). Although grasshoppers were fed *ad libitum* in our experiment, more individuals in a cage consume more and may feed on the more nutritious parts of leaf blades first. Hence, food quality may have been indirectly affected in a way that high population density could reflect poor conditions. However, our analysis showed that survival within cages did not predict wing development as well as nymphal density did, indicating that the population density itself is more likely the proximate cause of long wing development rather than poor conditions.

If population density is the main cause for the development of long wings, the actual mechanistic trigger may be visual, tactile, or olfactory cues (Anstey et al., 2009; Rogers et al., 2003; Heifetz et al., 1996). Visual cues do not seem to play a role in phase transition of locusts (Heifetz et al., 1996) and little is known for other Orthoptera. In contrast, there is strong evidence from phase-polyphenic locusts (phase polyphenism is the differential expression by the same genotype of morphological, physiological, and behavioural traits when being solitarious or gregarious) that tactile stimuli induce the development of the gregarious phase (Lees, 1967; Johnson, 1965). We also consider tactile cues to be the most likely cause in our study. There is also some evidence for olfactory effects in locusts, although effects seem smaller than for tactile influences (Heifetz et al., 1996). Since we found low-density cages with short-winged individuals located right next to high-density cages with long-winged individuals on many occasions, we consider volatiles to be a less likely cue in our case. However, variance decomposition shows about 8% of cage-related variation, some of which may include non-tactile interactions among cages. It remains open whether the density affecting morph development is only the conspecific density. Since many grasshopper species are ecologically, morphologically and physiologically similar and co-occur in communities (Liu et al., 2007; Gardiner and Hill, 2006; Gardiner et al., 2002; Belovsky and Slade, 1995; Ritchie and Tilman, 1992), we do expect that total grasshopper density – not only conspecifics – causes the development of long wings. To our knowledge, such interspecific effects have not been tested.

It may be argued that wing-dimorphism does not directly map to actual dispersal if, for instance, long-winged individuals still have reduced muscular structures. However, there is evidence from our study species, the meadow grasshopper, that long-winged individuals are actually dispersive. At high altitudes in the Harz mountains (Germany) there are no stable meadow grasshopper populations, but in some years long-winged individuals occurred in some numbers (Meineke, 1994). These individuals likely came from populations at lower altitudes, even though those populations were predominantly short-winged (Meineke, 1994).

The development of long wings did not show any clear density-dependent switch point in our study. For the phase-polyphenic locust *Schistocerca gregaria* (Orthoptera, Acrididae), there seems to be a more pronounced density threshold between phases (Heifetz et al., 1996). We hypothesize that the need for swarming in locusts calls for a harmonized density threshold – otherwise only small swarms would be formed at medium density. In non-swarming species, such as the meadow grasshopper, the decision to develop long wings seems to be taken at a more individual level. The phase polyphenism in locusts involves a whole suite of traits and therefore reflects a much more complex phenomenon than the wing dimorphism we studied. Very few species worldwide are phase-polyphenic (Song, 2011), which makes the wing dimorphism of the meadow grasshopper more representative of most grasshoppers. Our data suggests that significant, but fuzzy density-dependence may apply to those more widespread wing dimorphisms.

Once wings are developed in grasshoppers, their length is fixed for life. The determination of which wing morph an individual will develop has to occur during ontogeny. The restricted developmental periods of sensitivity to the environment determining the developmental trajectory are called sensitive phases (Sachser et al., 2020; Zera and Denno, 1997). Although our design was not optimized to test for specific sensitive phases, our data suggest that long-wing development depends mostly on densities during mid to late nymphal stages (*nymphal density* in our analyses) rather than on early-stage densities (*hatchling density*). This appears ecologically plausible, because most mortality occurred during the earliest stages of development, making the density of advanced nymphs more ecologically relevant with respect to overall competition throughout life.

We found that the production of long-winged individuals was family-dependent: some families produced more long-winged offspring at comparable density, others fewer, and this pattern was confirmed by the animal model analysis. This suggests that there is (polygenic) genetic variation for a threshold trait in the population. However, there is the alternative (or additional) option that epigenetic variation contributes to the development of long wings. In our two-generation pedigree, it is not possible to distinguish between genetic and epigenetic transmission. However, the parental densities were not manipulated (and putatively random with respect to offspring nymphal densities), such that intergenerational epigenetic transmission is not specifically indicated. In any case, the basis seems to be polygenic rather than monogenic (whether based on sequence variation or epigenetic modification), since monogenetic inheritance would have produced stronger patterns in our data – even if mixed penetrance of genetic variants may blur the pattern.

The development of long wings has important implications for how individuals can improve the phenotype-environment match. Two important mechanisms that allow individuals to adjust the phenotype-environment match are niche conformance – the adjustment of the phenotype to match the environment – and niche choice – the selection of habitat patches that best match the phenotype (Trappes et al., 2022). Induced long wings represent a form of niche conformance, since they are a response to local crowding. At the same time, long-winged individuals (whether induced or genetically determined) can sample a wider range of environments, which has the potential to improve the phenotype-environment match. Wing dimorphism, therefore, has dual implications for the formation of individualized ecological niches. Our data show that phenotypic plasticity and heritable variation are both involved in wing morph determination in grasshoppers, implying that niche conformance and niche choice can evolve in natural populations.

Overall, 48.5% of the phenotypic variance was explained by known environment (density, 22.4 %), additive genetic (18.9 %), and shared early rearing effects (7.2 %). While this might seem low at first glance, it is quite high for binary traits that inherently have a large residual component in polycausal effects. Additionally, we showed that environment and genes interact in the wing dimorphism of the meadow grasshopper, as opposed to what was shown for the brown and whitebacked planthoppers (Matsumura, 1996; Zhang et al., 2023). These contrasting results suggest that the evolutionary implications of wing dimorphism may differ from one taxon to another, and emphasize the importance of jointly examining the genetic and environmental factors of wing dimorphism in future studies.

Our data shows that wing determination in the meadow grasshopper is heritable and seems to be controlled by a polygenic complex coding for a threshold response to population density experienced during advanced nymphal stages. Population density and genetics explained a substantial proportion in wing morph determination. This suggests niche conformance traits can evolve through natural selection. Furthermore, our results highlight that studies on wing morph determination should consider these two factors jointly rather than separately in order to draw conclusions that are translatable to field populations.

## Acknowledgments

We thank the team of helpers for attentively maintaining the grasshopper population and in particular Ilka Wolf for coordinating the animal husbandry work. This work was supported by the Deutsche Forschungsgemeinschaft (DFG) (316099922, TRR 212, to H.S.) and an IMPRS stipend from the Max Planck Society (to M.V.).

## Author contributions

Cabon and Schielzeth conceived the ideas and designed methodology; Varma, Ebeling and Schielzeth collected the data; Winter, Cabon and Schielzeth analysed the data; Schielzeth acquired the fundings; Cabon and Schielzeth led the writing of the manuscript. All authors contributed critically to the drafts and gave final approval for publication.

## Data and code availability

Data and code will be uploaded on Datadryad upon acceptance of the manuscript.

